# Computer-assisted beat-pattern analysis and the flagellar waveforms of bovine spermatozoa

**DOI:** 10.1101/2020.03.06.972646

**Authors:** Benjamin J. Walker, Shiva Phuyal, Kenta Ishimoto, Chih-Kuan Tung, Eamonn A. Gaffney

## Abstract

Plagued by hurdles in information extraction, handling, and processing, computer-assisted sperm analysis (CASA) systems have typically neglected the complex flagellar waveforms of spermatozoa, despite their defining role in cell motility. Recent developments in imaging techniques and data processing have produced significantly-improved methods of waveform digitisation. Here, we utilise these improvements to demonstrate that near-complete flagellar capture is realisable on the scale of hundreds of cells, and, further, that meaningful statistical comparisons of flagellar waveforms may be readily performed with widely-available tools. Representing the advent of high-fidelity computer-assisted beat-pattern analysis (CABA), we show how such a statistical approach can distinguish between samples using complex flagellar beating patterns rather than crude summary statistics. Dimensionality-reduction techniques applied to entire samples also reveal qualitatively-distinct components of the beat, and a novel data-driven methodology for the generation of representative synthetic waveform data is proposed.

## 1 Introduction

In the context of analysing spermatozoa, information handling and processing has, in general, substantially lagged behind advances in microscopy. For instance, phase contrast microscopy was developed and commercialised in the 1930s and 1940s [1] and enables a straightforward visualisation of the spermatozoan flagellum, but the prospect of capturing and utilising the wealth of flagellar information beyond hand tracings [2] was not feasible until the 1980s, with software packages such as ‘BohBoh’ introducing a degree of automation [3].

However, the extensive user intervention still required for such packages as well as basic thresholding based techniques, for example by Smith et al. [4] who reported on the beats of 36 swimmers, has to date largely restricted population-level computer-assisted sperm analysis (CASA) to head-based investigations [5–8] and assessments of flagellar morphology [9], with flagellar waveforms neglected. Despite such limitations, and its only partial acceptance in clinical diagnostics [10], CASA has also found extensive application in reproductive toxicology, quality assurance for semen marketing in livestock breeding, improvements in sperm technologies such as cryopreservation, and studies of basic sperm function [11–13].

Furthermore, digital videomicroscopy that captures flagellar movement potentially contains a wealth of additional information, motivating the need for more user-friendly, less user-intensive techniques to capture the moving flagellum. A recent tool has been developed for this purpose with phase contrast microscopy by Gallagher et al. [14], though it is limited due to its current inability to capture the distal region of the flagellum, despite the reported importance of the distal flagellum in motility mechanics [15]. Other recent works have similarly developed techniques to capture the beating flagellum, including the distal region, most notably the three-dimensional capture of the flagellar waveform achieved by Daloglu et al. [16] with holographic imaging. However, compared to phase contrast microscopy, this is a highly-specialist imaging modality requiring custom circuitry together with sophisticated hardware and processing, whilst the former is already ubiquitous in theriogenology and andrology facilities [6, 9], and also in the study of a wealth of eukaryotic monoflagellates [17].

Proposing a different basis for flagellar capture, the automated methodology of Walker et al. [18] utilises the approximately-consistent width of the flagellum to distinguish it from the head of the swimmer, not relying on explicit image contrast between cell components and thus potentially applicable to typical videomicroscopy even with low signal-to-noise ratios, with far less user-processing than required with many systems. Thus, as an initial aim of this work, we seek to make available a large dataset of near-complete flagellar waveforms for hundreds of motile bovine sperm cells via the methodology of Walker et al. [18], highlighting that the fundamentals of a new generation of *computer-assisted beat-pattern analysis* (CABA) for essentially the whole and motile flagellum is realisable for eukaryotic monoflagellates on current, widely-available hardware utilising simple, intuitive image processing.

Whilst the ability to readily document the flagellar beat in high fidelity using commonly-available imaging modalities has significantly broadened the potential scope of CASA, arguably of greater importance is the prospect of detailed quantitative analysis incorporating the entirety of the motile flagellar waveform. Such statistical analyses, with the power afforded by non-small sample sizes, would provide the capabilities required for a new generation of swimmer evaluation studies, based on quantitative assessment of not only coarse summary statistics, as is the current standard [6, 8], but also the complex time-evolving shape of the flagellar beat. However, even given detailed waveform data, there is no clear or established methodology for performing statistical comparisons and hypothesis testing on flagellum data.

In numerous previous works [19–23], the waveforms of beating flagella have been simplified via dimensionality-reduction techniques, a pertinent example being the principal component analysis (PCA) employed by Ma et al. [21] though not in a statistical context, with only seven swimmers analysed. This standard technique aims to decompose a dataset into a small number of representative modes, capturing the variance of the underlying observations. Here, building upon captured waveform data, our further aim will be to use such dimensionality-reduction techniques, but applied to entire samples, to generate distributions that describe the flagellar waveform and its variation across a sperm population. The fundamental objective of this work will then be to use these population-level distributions associated with flagellar waveforms to enable flagellar-based sample comparisons via the subsequent application of common techniques for statistical hypothesis testing. In particular, these techniques will be illustrated by considering the beating patterns of post-thaw spermatozoa that present with distal cytoplasmic blebbing, assessing whether or not their waveforms may be distinguished from those unaccompanied by signs of damage. In turn, this showcases how CABA may be used to quantitatively compare motile spermatozoan flagella across populations.

Finally, a further objective will be to utilise the captured waveform data in the generation of evidence-based synthetic flagellar waveforms by a simple sampling method. In doing so we will provide a methodology for the construction of arbitrary numbers of evidence-based synthetic flagellar waveforms for use in biophysical population modelling, with current modelling frameworks accommodating hundreds [24] and even up to one thousand [25] virtual spermatozoa. In turn, this will enable future studies to assess the physical consequences of evidence-based population variability in explorations of flagellated swimmer active matter.

In summary, we will generate and present a large dataset comprising of the fine details of swimmer waveforms from phase contrast videomicroscopy, utilising widely-available non-specialised hardware and simple, effective image processing. We will then leverage this dataset to produce synthetic waveform data, proposing a novel method of beat pattern generation that captures key features of the spermatozoan beat, envisaging its use in future biophysical modelling applications. Equipped with rich kinematic data, we attempt to decompose the flagellar beat into its most prominent components, highlighting the potential for high-fidelity flagellum datasets such as that presented here to facilitate biological enquiry at population level into spermatozoan flagellar dynamics. Further, we will demonstrate by example that detailed qualitative comparison of samples based on highly-resolved waveform information is a realisable direction for CASA and represents a new generation of automated flagellum analysis, making use of dimensionality reduction techniques and standard statistical testing.

## 2 Results

### 2.1 Large scale quantification of flagellar beating

#### 2.1.1 Waveforms, paths, and periods

We imaged the beat patterns of swimming bovine spermatozoa in 1% methylcellulose dissolved in Tyrode Albumin Lactate Pyruvate (TALP) medium, and digitally captured approximately 90% of the planar-beating flagellum. This dataset is freely available (as detailed in the data access statement), and presents spatially and temporally-smoothed parameterisations of the flagellar waveforms for 216 swimmers, given over a single period, aligned in phase, and normalised as described in the Methods and Materials. We present a sample frame, identified flagellum and a captured beat pattern in fig. 1. Our dataset represents the digitisation of flagellar waveforms not previously captured on this scale via the common imaging modality of phase contrast, as previous distal capture is limited at approximately 70%[14]. We present this dataset of automatically captured and smoothed flagellar beats in both the Cartesian and tangent angle forms (see Methods and Materials), accompanied by the computed beating period and truncated flagellum length. We also provide raw tracking data generated by the software TrackMate [26], which may be used in isolation to compute the wide range of existing CASA measures [8].

**Fig 1.**
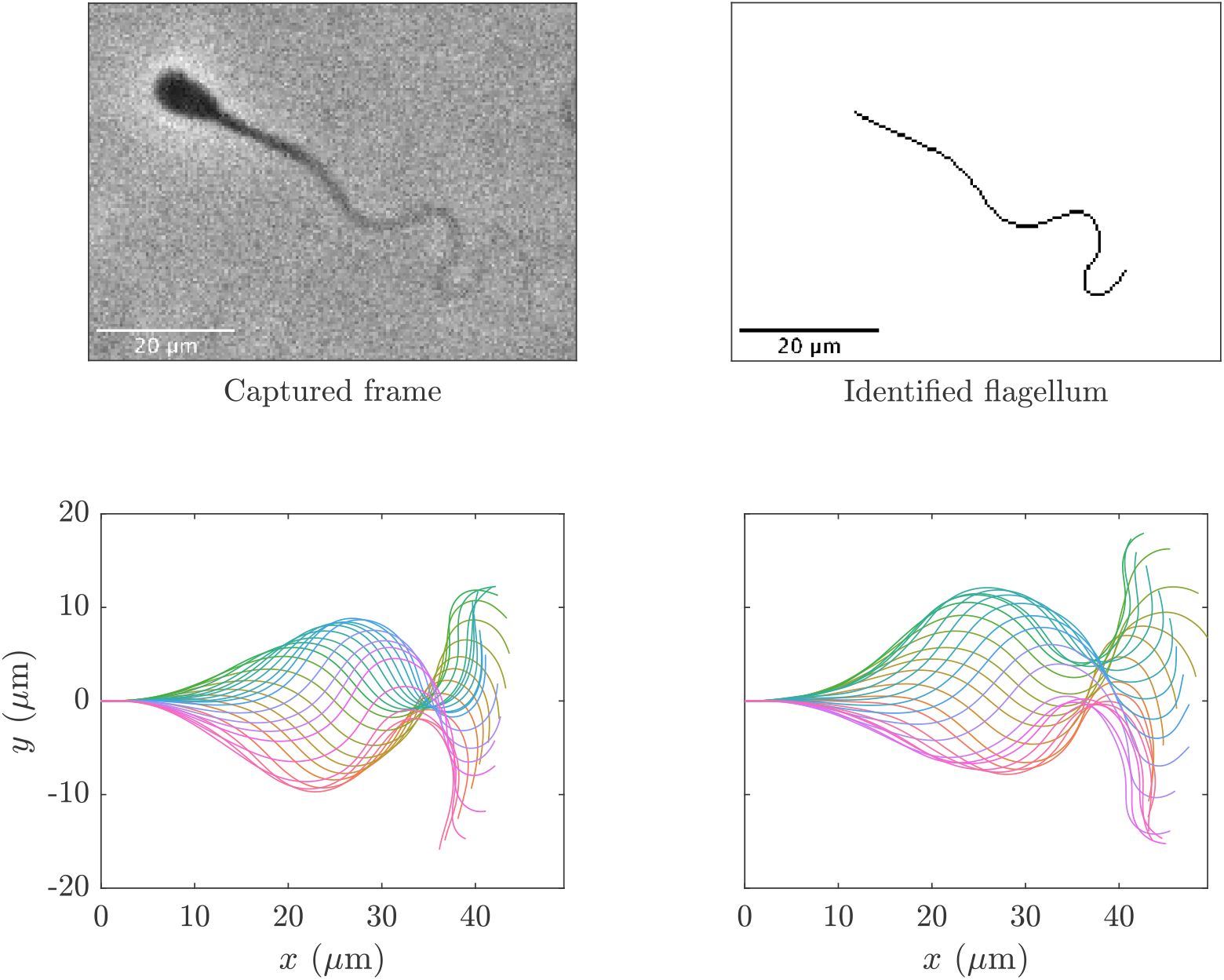
A sample captured frame from a fresh spermatozoon, the identified flagellum, and a complete waveform, accompanied by a sample synthetic waveform constructed from the presented dataset utilising the proposed method of generation with *N_s_* = 10. The upper panels show the raw image data alongside the flagellum as identified using the automated methodology of [18], with the vast majority of the flagellum being captured. The lower left panel displays a captured waveform over a single beating period, with the sample synthetic beat shown in the lower right panel having been generated following eq. (1), drawing flagellar arclength and beating period from their respective empirical distributions. In these lower panels, plotted in different colours are the shapes of the flagella at select timepoints within one period, with the base of each flagellum situated at the origin of their respective Cartesian coordinate system. We remark that these waveforms are qualitatively similar, with the proposed sampling method replicating the mean and variance of the empirical waveform distribution by construction. The waveforms in the lower left and right panels have been scaled up by the original and sampled flagellum lengths in order to enable comparison to imaging data.

#### 2.1.2 Sampling waveforms

Utilising this rich dataset of captured spermatozoa beats, we propose a novel method for the generation of synthetic waveforms that follow the approximate distribution of digitised waveforms. Herein, *s* is an arclength parameter that has been rescaled for each individual swimmer by the flagellum length, so that *s* ranges from zero to one. Similarly, *t* denotes dimensionless time over a single beating period, with time having been normalised for each swimmer by their beating period so that here *t* also lies between zero and one (see Methods and Materials for full details). Working in the framework of angle parameterisations, fixing some integer *N_s_*, and uniformly sampling *N_s_* waveforms *θ*_1_, …, *θ_N_s__* from our entire dataset, we can form a synthetic angle parameterisation *S* via

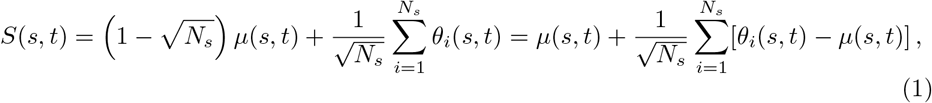

parameterised by the normalised arclength *s* and rescaled time *t*, where *μ*(*s, t*) denotes the average of the angle parameterised waveforms in our dataset. Given a synthetic angle parameterisation, we generate a waveform by uniformly sampling an arclength and period from the constructed dataset, together constituting a full description of a generated flagellar beat pattern. Synthetic waveforms created in this way retain the mean and variance of the original dataset, and hence approximate a representative spermatozoan waveform given approximate normality in the deviation of waveform angles from their mean. The latter assumption is informed by the distributions of PCA coefficients [27, Chapter 2], shown in the lower panels of fig. 4. We include in the Supplementary Material accompanying this publication an exemplar MATLAB^®^ code for generating synthetic waveforms in this manner, with a sample synthetic beat being shown in the lower right panel of fig. 1. It should be noted that, whilst here we sample from our entire waveform dataset, application of this methodology to stratified datasets may necessitate sampling only from subsets of the data.

### 2.2 Computer-assisted waveform analysis

#### 2.2.1 Refined, automated, and accessible flagellar analysis

In producing such a dataset of flagellar beating we have demonstrated that a new generation of computer-assisted beat-pattern analysis, CABA, is feasible and realisable with non-specialised hardware and software setups, having captured the beating patterns of hundreds of individual swimmers including a vast proportion of the flagellum using accessible image capture and processing. Such flagellar detail may enable the development of new measures of spermatozoan motility and quality, for example with reliable estimates of swimming efficiency and power output now possible.

An additional pertinent metric for the assessment of spermatozoa may be the maximal tip curvature, defined in the Methods and Materials, which was previously difficult to obtain due to limitations of flagellum tracking and imaging. The importance of such a measure, which quantifies the magnitude of bending in the distal flagellar region, is suggested in the recent work of Neal et al. [15], wherein changes in the curvature of the distal tip of model flagella are reportedly correlated with changes in efficiency and swimming speed. This suggestion is simply one of a multitude of measures that one may take from near-complete flagellar data as is presented here. Indeed, with the future incorporation of elastohydrodynamical models of flagellar beating, one may even be able to quantify in unprecedented detail mechanical aspects of the flagellum, for instance the net activity of molecular motors within the axoneme, to provide a novel and in-depth perspective on swimmer activity with potential links to individual viability [28].

#### 2.2.2 Contributions to the flagellar beat

Performing principal component analysis (PCA) *on the entire population of swimmers* allows us to quantify the components of the flagellar beating of bovine spermatozoa, accounting for variation throughout the individual beats and across the population as a whole. In fig. 2 we present depictions of the contributions of the first three Cartesian PCA modes of the waveforms, with these modes corresponding to 75%, 7.5% and 6.9% of the total variance, respectively. The first PCA mode can be seen to generate the most significant component of the flagellar beat, an actuation perpendicular to the midline of the waveform. The second mode contributes compression and extension of the flagellum in the direction of the midline, also introducing a slight bend. Most recognisable in terms of its visual contribution to the waveform, the third mode introduces the characteristic sinusoidal shape to the beat, with the amplitude appearing to increase with the distance from the flagellum base.

Not only are the effects of the modes visually distinct, but their contributions differ over time in both magnitude and phase. Shown also in fig. 2 are the average values of the first three PCA coefficients over a beating period, and we note that their magnitudes are comparable as the modes have been, without loss of generality, suitably normalised. The first mode, corresponding to 75% of the variance, is of much greater magnitude than the less significant components, thus it is the dominant contribution to the amplitude of the overall beat. Representing lesser contributions to both the variation and the beat amplitude, the coefficients of the higher order modes are additionally out of phase, both with each other and the dominant mode. With the third mode, generating the stereotypical sinusoid-like waveform, for example, this phase difference implies that the characteristic shape of the beat lags behind the main oscillatory component, at least in this average waveform.

**Fig 2.**
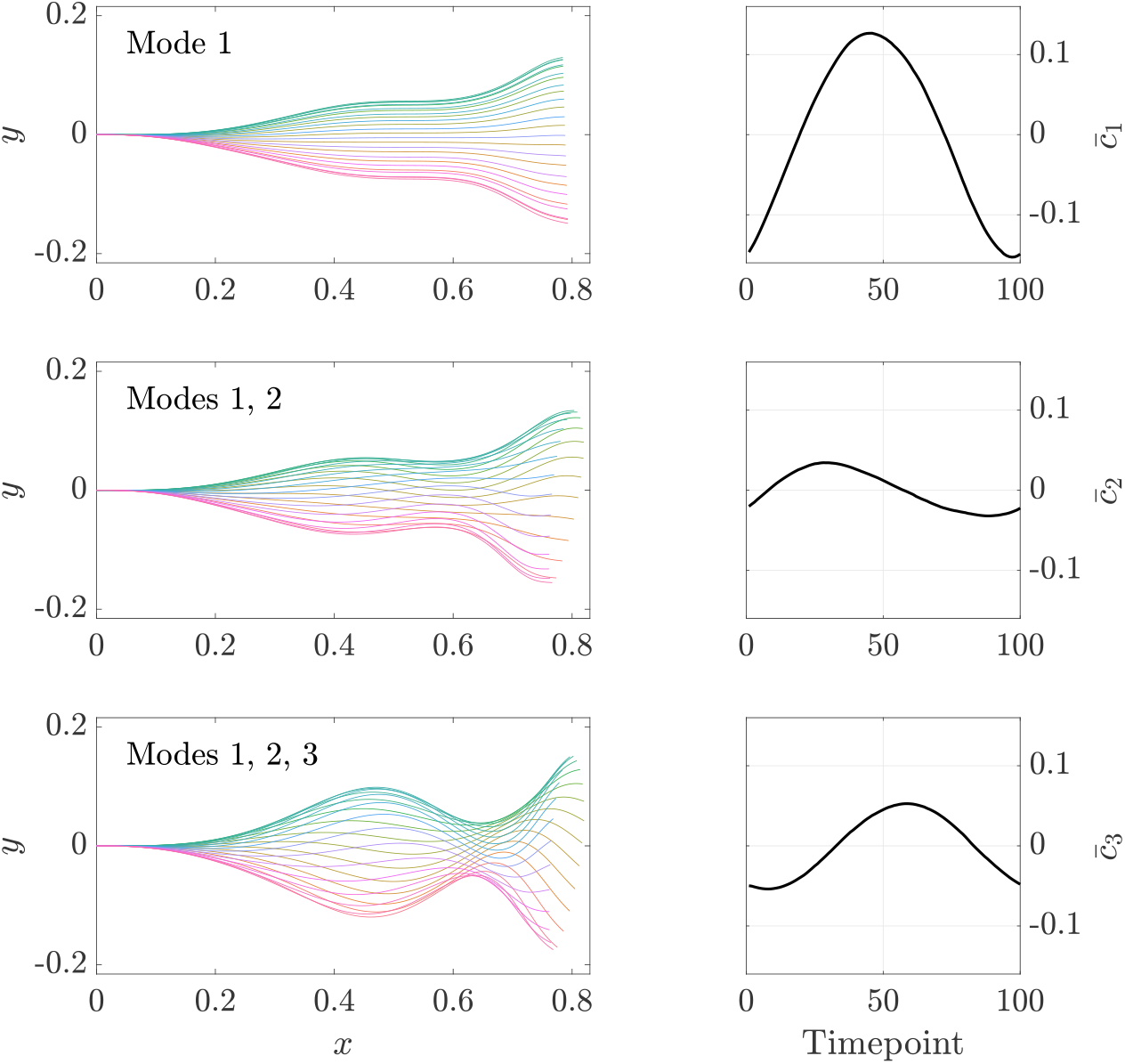
The cumulative contributions of the most prominent Cartesian PCA modes, added to the population mean. The left panels showcase the effects of the three most-significant modes, presented as the normalised flagellar waveforms generated by the modes along with the average Cartesian PCA coefficients, shown over a single beating period and with the base located at the origin. These average PCA coefficients are denoted by 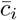 for *i* = 1, 2, 3, with their evolution shown over time in the right panels. We can see that the dominant mode, capturing 75% of the population variance over a beating period, prescribes the amplitude and overall shape of the flagellar beat, with the maximum of 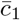 greater in magnitude than the other coefficients. The contributions of the higher order modes are out of phase with the primary mode and of reduced magnitude, though we note that the third mode appears to generate the recognisable sinusoidal shape that is characteristic of many bovine spermatozoan beat patterns. Different colours correspond to different timepoints throughout the flagellar beat, consistent between rows. Without loss of generality, all modes and coefficients have been normalised to enable direct comparisons between the magnitudes of PCA coefficients.

Repeating the above principal component analysis with waveforms represented as angle parameterisations yields analogous results, though with different principal components owing to the nonlinear transformation that takes spatial data to angle form. It is not clear that these components can be separated into the constituent parts of the flagellar beat identified by analysis of the spatial data, highlighting that in general it may be of benefit to consider waveforms not only as reduced angle representations but also in spatial form, with the differing approaches potentially yielding different insights into the flagellar beat.

### 2.3 Variation across spermatozoa

With hundreds of spermatozoa imaged and their flagellar waveforms digitised, we perform several exemplar comparisons of the flagellar beat between two different samples, referred to herein as sample A (fresh) and sample B (frozen), imaged with two different cameras (see Methods and Materials). This highlights that meaningful comparison between swimmers is not simply limited to coarse summary statistics and head tracking, but may take into account the fine details of the flagellar waveform. Inumerable additional comparisons are possible in future studies given the presented dataset and the CABA methodological framework, with sample sizes providing statistical power and flagellum detail enabling the exploration of novel beat metrics.

#### 2.3.1 Beating period

A fundamental feature of any periodic beating is the extent of its period. We define the period as the time interval after which the flagellum most closely resembles its initial configuration, matching our intuitive notion of a beating period, as detailed in the Methods and Materials, and circumventing cumbersome and problematic definitions via Fourier spectra. In fig. 3a we report the period of the flagellar beat for each of the individuals in samples A (fresh) and B (frozen), with the medians shown as dashed lines. The two distributions appear starkly distinct to the eye, supported by a Wilcoxon rank-sum test at the 1% significance level, in addition to a significant Kolmogorov-Smirnov test statistic. Hence there is sufficient evidence to suggest differences between the samples, with sample A having approximately twice the beating period of sample B on average.

**Fig 3.**
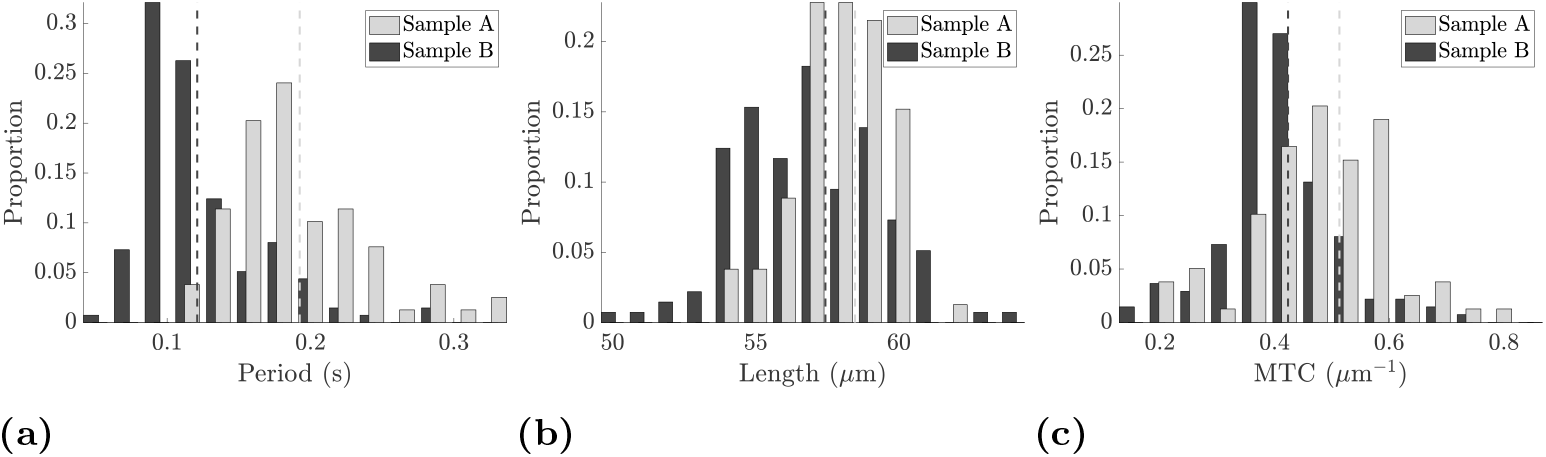
Empirical distributions of beating period, flagellum length, and maximum tip curvature (MTC), shown as histograms and separated into sample A (fresh) and sample B (frozen). (a) Distributions of the period of the flagellar beat, from which we can visually identify differences between sample A and sample B. Though less pronounced, significant differences between both medians and entire distributions can also be seen for the observed flagellum length and the maximum tip curvature, shown in (b) and (c), respectively. Thus, via these simple exemplar metrics, we are able to identify differences between samples of spermatozoa. The medians of each distribution are shown as dashed vertical lines, with distributions corresponding to sample B shown darker, and all differences in distribution reported here are confirmed via Kolomogorov-Smirnov tests at the 5% significance level, with medians similarly assessed by the Wilcoxon rank-sum test.

#### 2.3.2 Flagellum length

fig. 3b shows the distribution of flagellum length for each sample. We again see statistically significant differences between the distributions, in both median and overall distribution. In line with noted morphological homogeneity in bovine spermatozoa [29], apparent from the distributions of arclength is the low variance about the mean, true of both samples individually and additionally when combined (not shown). Such tight spread supports our rescaling of all flagella by total length, enabling later comparison by principal component analysis.

#### 2.3.3 Extremes of distal bending

With such extensive flagellar detail captured, we can begin to consider details of the waveform previously unavailable on this scale, in particular, the maximal distal curvature. Suggested by [15] to be correlated with swimming efficiency, we compute the maximal tip curvature (MTC) of each of the swimmers throughout their beat, shown as histograms in fig. 3c stratified by sample. Again we observe statistically significant differences between the two samples, consistent with the PCA analysis that follows below and highlighting how fine details of the flagellar beat can be used to identify differences between populations of swimmers in terms of suggested surrogates of functional performance, such as swimming efficiency.

#### 2.3.4 Waveform

Having performed principal component analysis (PCA) on the entire population as a whole in Cartesian form, we can interrogate the resulting coefficients to identify any significant differences in the beating patterns of samples A and B. In fig. 4 we present the distribution of the coefficient of the first Cartesian PCA mode for each individual normalised waveform, separated by sample and shown over the course of a single beat. Shown in the upper panels are the time series of the first PCA coefficient, *c*_1_, for each swimmer, accompanied by the evolution of the sample means plus and minus the standard deviation (shown as darker, heavier curves). To the resolution shown in the figure, sample B appears to display larger variance during the central 40 timepoints of the beating than sample A, in addition to an increased mean value. This is more apparent in the lower panels, which for each timepoint show the coefficient distribution as a colour-coded histogram, where higher values of the empirical probability distribution are shown as darker regions. The higher variance of sample B can be observed, in addition to the consistently-unimodal nature of the distribution of coefficients over swimmers.

**Fig 4.**
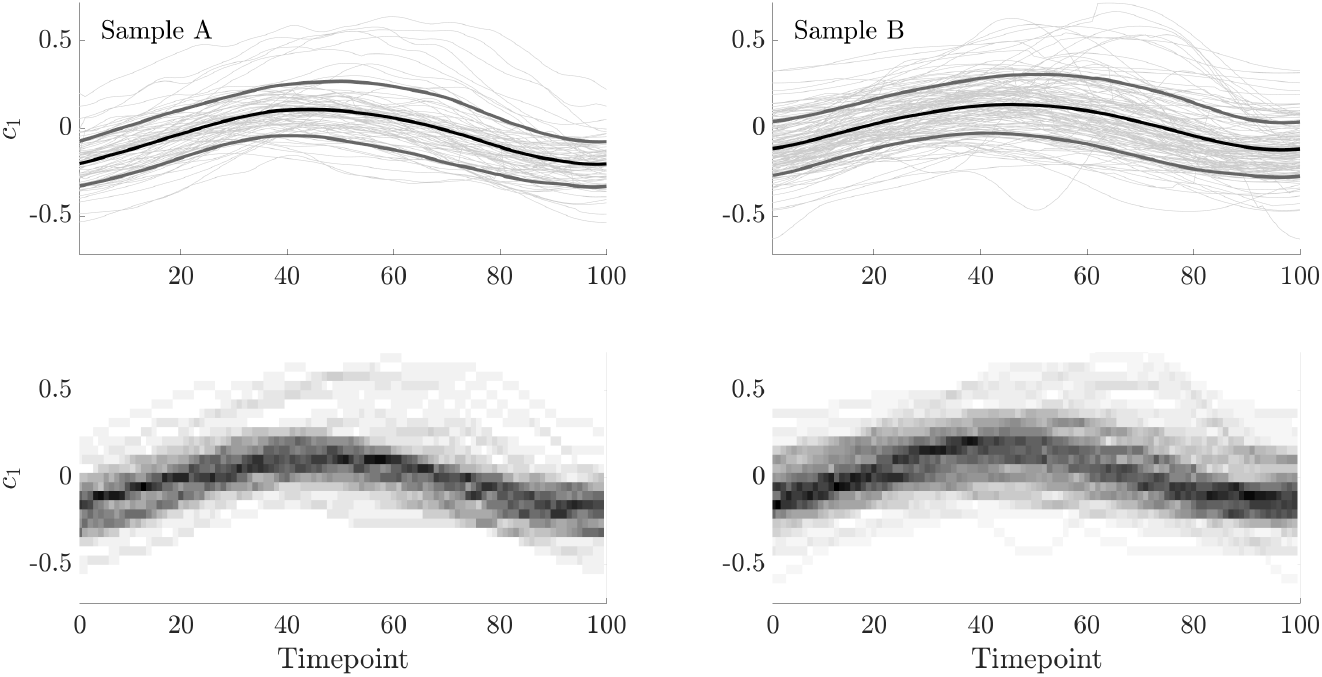
Distributions of the first Cartesian PCA coefficient, *c*_1_, over the course of a single beat, with samples A and B shown in the left and right panels, respectively. In the upper panels, we show the traces of *c*_1_ for each swimmer as light grey curves, accompanied by the sample mean, presented as a black, heavy curve. Shown also is the mean plus or minus a single standard deviation, here drawn as heavy grey curves flanking the mean. In the lower panels, we show the empirical distributions of the first PCA coefficient, constructed by computing a histogram at each timepoint and associating each bin probability with a greyscale value that is uniformly varying on the interval [0, 1]. Darker regions correspond to high-density regions of the histograms, with the modal values approximately following the mean curves shown in the top panels, consistent with approximate normality. Visual comparison between samples is difficult, though increased variance appears to be present in sample B compared to sample A between timepoints 40 and 80. In both samples we recognise an approximate sinusoidal shape of the mean and mode, suggesting that a simple sinusoidal fit may be appropriate to capture the evolution of the first PCA coefficient over time.

Further to qualitative comments that speculate differences between the two samples, we may also conduct quantitative comparisons. Owing to the unimodality of the distributions and the simple temporal form of the means of the first PCA coefficient, we may compare the fits of a linear model for PCA mode coefficients via the distributions of the linear fitting parameters in order to examine statistical differences between samples. With the form and details of the linear model presented in the Methods and Materials, in summary we find that simple sinusoidal fits to the dataset reveal a statistically significant difference between the two samples, in concurrence with the difference in means that can be seen between the left and right panels of fig. 4, though such a difference is hard to detect, and especially quantify, by eye. In fig. 5 we explicitly show empirical histograms for the parameters *α* and *β* in a statistical fit of the first Cartesian PCA coefficient, *c*_1_, to a sinusoidal function,

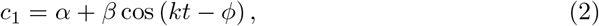

where wavenumber and phase parameters *k* and *ϕ* are first approximated by non-linear least squares fitting of the mean coefficient, 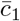 (See fig. 2, and the section entitled “Statistical tests” within Methods and Materials for further details). In particular, such tests facilitate detecting and testing for differences between sample A (fresh) and sample B (frozen) more readily. Thus, via consideration of the details of the flagellar waveform, subtle but significant differences between samples may be both automatically identified and rigorously quantified, utilising an approach that may be readily extended to include additional PCA coefficients and alternative methods of dimensionality reduction.

**Fig 5.**
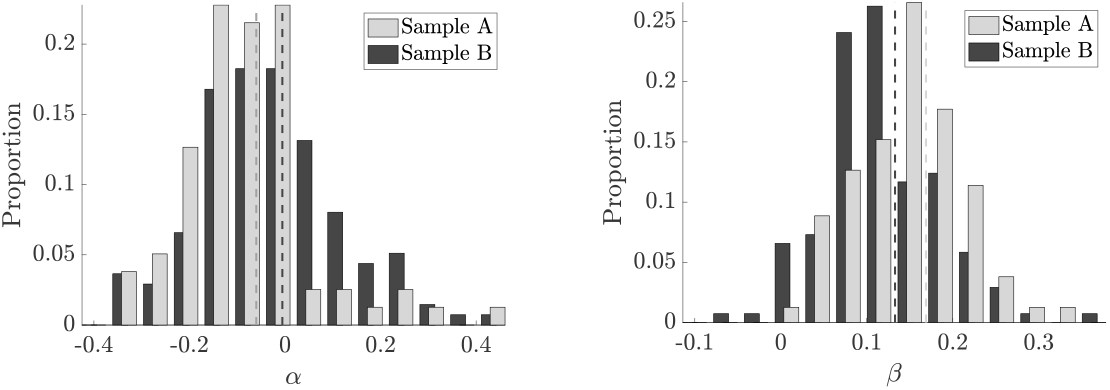
Marginal empirical distributions of fitted parameters (see eq. (2)), corresponding to sinusoidal fits for the first PCA coefficients of each swimmer, stratified by sample. Having fit the coefficients of a simple sinusoidal model to *c*_1_, as suggested by fig. 4, we show the empirical distributions of the fitted parameters as histograms, with those corresponding to sample B shown darkest. Visual differences between samples in median (shown as dashed curves) and variance are evident by eye for each of the fitted parameters, supported by Kolmogorov-Smirnov tests at the 5% significance level, noting that there is no assumption about the underlying distributions in this test. Thus, we can quantitatively distinguish between the flagellar beating of two samples of spermatozoa.

### 2.4 Assessing impacts of distal cytoplasmic blebbing

With the spermatozoa in sample B having undergone a freezing and thawing process, we note that a significant proportion (25%) of these motile individuals exhibited visible signs of damage, specifically distal cytoplasmic blebbing. We show two example such swimmers in fig. 6 and henceforth refer to swimmers with visible distal cytoplasmic blebbing as *blebbed*, with visually unaffected individuals being referred to as *unblebbed*. Having manually identified blebbed cells, we repeat the comparative analysis performed above for samples A and B on the motile blebbed and unblebbed subsamples.

**Fig 6.**
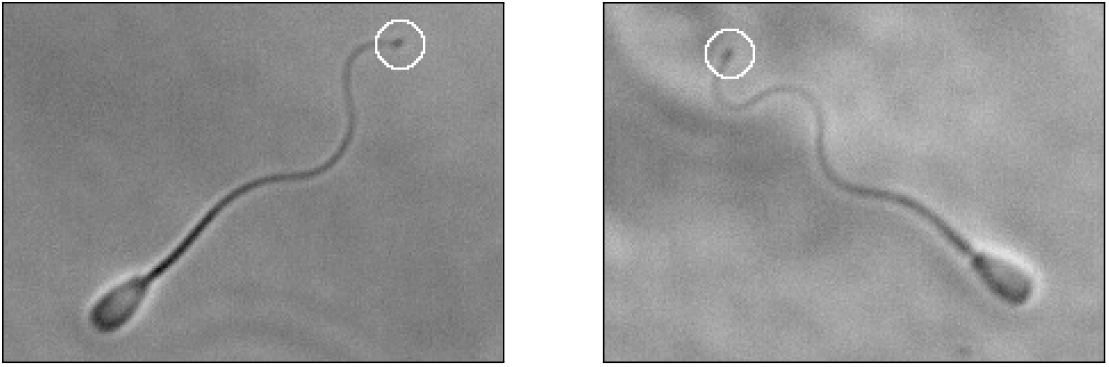
Visible signs of flagellar damage, with select frames taken from sample B (frozen). We see evidence of distal cytoplasmic blebbing, present here as darker, broader regions towards the tip of the flagellum of affected swimmers, highlighted by an annotated white circle. In our sample of 137 swimmers that had undergone a freezing and thawing process, approximately one in four individual spermatozoa was found to exhibit visually identifiable blebbing.

We find that there is a statistically significant difference in the distributions of flagellum length between blebbed and unblebbed swimmers via a Kolmogorov-Smirnov test at the 0.5% significance level, although the same test is inconclusive for distributions of maximal distal curvature and beating period. The Wilcoxon rank-sum test applied to the beating period does however suggest a difference in medians at the 5% significance level between blebbed and unblebbed periods, with the blebbed swimmers possessing a lower median compared to unblebbed cells. Finally, and consistent with inconclusive results pertaining to maximal distal curvature, we find no statistically significant differences between the waveforms as a whole by considering linear model fits of the PCA coefficients. In turn, this statistically evidences that, despite the freeze-thaw transitions inducing distal flagellar blebbing in these bovine spermatozoa, the distal beating pattern of the blebbed spermatozoa is nonetheless preserved relative to those with no visible damage.

## 3 Discussion

In this work we have produced and analysed a dataset of the flagellar beats of over 200 bovine spermatozoa and have described a methodology and pipeline that is easily integrated with phase contrast microscopy, which is ubiquitously available in many theriogenological, andrological and cell motility laboratories. Whilst also of pertinence to a broad biological community, we anticipate that the data generated in this study (available as described in the accompanying data access statement) will also be of intense interest to the biophysical modelling community. Datasets generated with the same level of flagellar detail and scale have, to the best of our knowledge, been previously unavailable, with more of the distal flagellar tip being captured here than has been realised on large scales prior to this study, with the notable exception of the recent work of Daloglu et al. [16] that applied sophisticated holographic imaging to capture beats in three dimensions. Easily-obtainable waveform data with the level of fidelity presented in this study has the potential to further the vast research area of heterogeneous population modelling, allowing the emergence of the next generation of evidence-based biophysical modelling for cell populations, spermatozoan behaviours in complex microenvironments, and more generally active matter physics. For example, the novel ability detailed here to generate synthetic waveforms will enable theoretical advances in understanding how realistic population heterogeneity impacts on the collective behaviours of active matter.

The detailed digitisation of flagellar beating and the widespread availability of such data additionally provides novel opportunities for quantitative comparison of flagellar beats across individuals, samples, and, in principle, species. Applying the comparative methodology described in this work to our two samples, we were able to identify statistically significant differences between the waveforms of the swimmers in each of the fresh and frozen samples using linear model fit parameters for the primary principal component analysis (PCA) coefficient, and thus examining more than simply coarse summary statistics such as period or wavelength, the latter of these not being uniquely defined in general. Whilst the impacts of these quantitative differences on overall motility are not clear, differences in suggested surrogates of functional performance, such as maximal tip curvature as a proxy for swimming efficiency, can be readily detected even in this initial study. Furthermore, future hydrodynamical and elastohydrodynamical computational studies could determine the extent of the effects, if any, of these dissimilar beats on swimmer progression and their metabolic demands, as well as their interactions with the microenvironment, including neighbouring swimmers. This example speaks more generally to the potential for quantitative and far-reaching comparisons of monoflagellated eukaryotic swimmers and their mechanical function with the availability of detailed, high-volume flagellar data, demonstrated to be realisable.

Owing to the significant sample sizes that may be analysed via our presented methodology, intrasample variations and associated comparisons may also be explored, for instance cytoplasmic morphology and its correlates. In particular, cytoplasmic blebbing has been noted in freshwater fish sperm, for example sturgeon [30], and attributed to osmolar stress. Here we have also observed cytoplasmic blebbing of the distal tip for one in four sperm that has undergone cell freezing and thawing. Even with such a morphological dichotomy in the presence and absence of blebbing within the frozen sample (B), we were unable to differentiate between the spatial aspects of the waveforms exhibited by each subgroup of sample B, including the waveform distal features, despite having amply large sample sizes to facilitate statistical power. This provides evidence that the damage inducing distal blebbing and the distal blebbing itself have no discernible effects on the beating characteristics of motile swimmers for reconstituted frozen bovine spermatozoa, with the exception of potential effects on beating period that warrant further exploration.

More generally, these example comparisons serve to illustrate how the presented techniques can be applied to assess the impacts, if any, of differences between swimmers on their overall motility function, utilising the details of highly-resolved flagellar waveforms. We have demonstrated that such analysis, performed here automatically on large sample sizes totalling hundreds of swimmers, and incorporating information from 90% of the length of the flagellum, is realisable with readily-available imaging and computational techniques, and thus represents a viable framework for a new generation of detailed flagellar CASA, and more generally the advent of computer-assisted beat-pattern analysis (CABA). The introduction of traditional CASA has had a substantial impact in theriogenology, for example enabling systematic quality control and assurance in the livestock breeding industry [9], as well as fundamental cell science, with extensive fundamental studies of sperm function [11–13]. Further, the expansion into near-complete planar flagellar waveform capture offers the prospect of novel directions for the exploration of male gamete motility, and flagellated or ciliated organisms more generally. The significance of high-fidelity analysis of flagella is further highlighted by the recent work of Neal et al. [15], with the overall efficiency of a spermatozoon suggested to be linked to distal curvature. Measurement of such curvatures requires the capture of large proportions of the flagellum, greater than previously available en masse with readily accessible microscopy though demonstrated here to be achievable on significant sample sizes. As an additional example, access to detailed flagellar data may also enable investigation into the symmetry of the spermatozoan beat, with significant asymmetry commonly associated with hyperactivation in mammalian spermatozoa but nevertheless exhibited at a much more subtle level here for bull, as illustrated in the lower left panel of fig. 1. Further, the method of quantitative analysis presented in this work need not be limited to the spermatozoa of any species, with the detailed analysis of beating flagella relevant to a range of organisms, from the biflagellated alga *Chlamydomonas reinhardtii* to the helically-driven bacterium *Escherichia coli*.

Datasets such as the one generated and presented in this work may additionally enable detailed investigation into the structure of the spermatozoan beat. In this study we have seen that the population-average flagellar beat can be deconstructed into distinct components, with the primary contribution generating the large amplitude motion perpendicular to the midline, and the formation of the stereotypical sinusoid-like shape lagging behind. With detailed flagellar waveforms now available on a large scale, similar analyses may lead to a deeper understanding of the flagellar beat, suggested here for bovine spermatozoa but potentially applicable in more generality. If used in combination with mechanical measures and elastohydrodynamic models of the flagellum, for instance, extensive in-depth datasets of in vitro waveforms may facilitate explorations of the mechanical regulation and actuation of flagella, enabling data-driven queries of the validity of popular hypotheses of flagellar control [28, 31–33].

In summary, we have investigated the beating characteristics of bovine spermatozoa, capturing more of the flagellar beat than previously acquired on such sample sizes without requiring sophisticated hardware or image analysis tools. With the resulting dataset publicly accessible, as described in the data access statement below, and with the prospect of readily producing further data, we have made available a wealth of kinematic data with potential for future use in in-depth quantitative studies that may lead to a deeper understanding of the flagellar beat, the biology of spermatozoa, and more generally active matter modelling. Indeed, such detailed waveform data will directly enable future biophysical modelling of heterogeneous swimmer populations, for example via the novel ability presented here to generate synthetic waveforms that retain key population characteristics. Most significantly, we have demonstrated that comparisons between the beating patterns of flagellated swimmers may be carried out quantitatively via nonstandard application of dimensionality reduction methods, taking into account aspects of the flagellar beat that have previously been inaccessible or neglected. In doing so we have showcased a realised pipeline for a new generation of computer-assisted flagellar analysis, readily achievable with current imaging and computational techniques. The presented approach leverages unprecedented flagellar fidelity with non-specialised imaging modalities to assess and classify the intricacies of the spermatozoan beat, facilitating potential future study into individual motility, viability, function and variability in motile flagellate populations.

## 4 Methods and Materials

### 4.1 Sample preparation & image capture

#### 4.1.1 Reagents & media

Tyrode Albumin Lactate Pyruvate (TALP) medium [34] was used as a standard medium for bovine sperm. TALP comprised of 3.1 mM KCl, 25 mM NaHCO_3_, 10 mM HEPES Free Acid, 99 mM NaCl, 0.39 mM of NaH_2_PO_4_, 25.4 mM of Na-lactate, 2 mM of CaCl_2_, 1.1 mM of MgCl_2_, 0.11 mg/mL Sodium pyruvate, 5μg/mL of gentamycin and 6 mg/mL of Bovine Serum Albumin. The final pH value of the TALP solution was 7.4. 1% (w/v) of Methyl Cellulose (MC, 4000 cP at 2%) solution was made by slowly stirring MC powder in TALP in room and chilled temperatures alternately. All fluids were equilibriated in a 38.5°C incubator (bovine body temperature) with 5% CO_2_ in humidified air before use. Rheology of the MC solutions were measured using a rotational shear rheometer (TA Instruments, DHR3) with a standard cup and DIN rotor at 38.5°C in oscillation mode. The measured moduli are shown in fig. 7. At the frequency of 10 Hz, the storage modulus, *G′*, and loss modulus, *G″*, were measured as *G′* = 0.73 Pa and *G″* = 5.37 Pa, respectively. As far as considering the dynamical frequency being less than 20 Hz, the rheological data were well approximated by the viscoelastic Maxwell model, from which we estimated the effective viscosity, 0.088 Pa·s, and the relaxation time, 0.0014 s.

**Fig 7.**
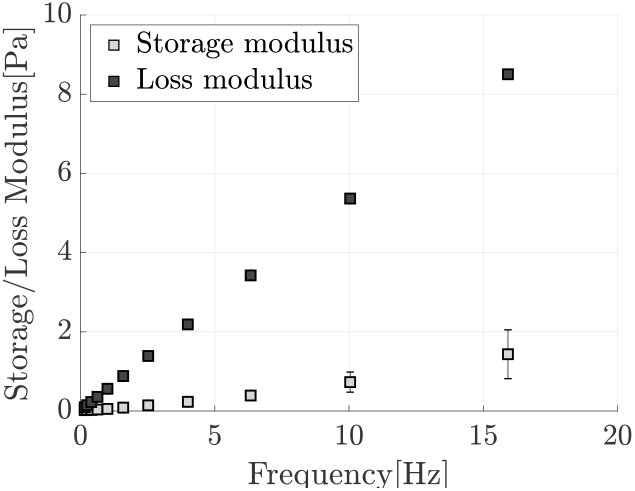
Measurements of storage and loss moduli for the methyl cellulose medium at various frequencies. We observe that both the storage and loss moduli, shown as light and dark squares respectively, are well-approximated by a linear viscoelastic Maxwell model. Error bars corresponding to the standard deviation of measurements are shown for each of the moduli, though only two such bars are visible at the resolution of this figure.

#### 4.1.2 Sample preparation

All bovine semen samples were kindly provided by Genex Cooperative, Inc. Sample A was a fresh semen sample from one bull, from which the beats of 79 sperm were captured. Shortly after semen collection, 1 mL of bovine semen sample was diluted in 5 mL of warm TALP, and then processed identically to Tung et al. [35]. Sample B was a frozen semen sample from another bull, from which the beats of 137 sperm were captured. Notably, as Samples A and B differ in both their handling and their source animal, we will not seek to draw biological conclusions from any further observed distinctions between them, and place emphasis wholly on the comparative methodology presented in this work that is exemplified by the consideration of these two samples. After collection, the semen was first diluted in OptiXcell extender (IMV t Technologies, L’Aigle, France), and then frozen according to the standard procedures followed at Genex [36]. The thawing process is similar to our previous procedures for thawing frozen semen straws [37]. The frozen semen was first thawed in a water bath at 37°C for 30 seconds, and then transferred on top of two layers (40% and 80%) of BoviPure diluted in BoviDilute. The live sperm were separated from the rest of the fluid by centrifugation (100 x g for 10 minutes), as they formed a pallet at the bottom of the tube. After removal of the supernatant, 3 mL of TALP was added and followed by another centrifugation (100 x g for 3 minutes) to wash the sperm pallet. 100 *μ*L of TALP was added to the sperm pallet to make the suspension, which was kept in an incubator maintained at 38.5°C with 5% of CO_2_.

#### 4.1.3 Image acquisition and experimental setup

Our previously developed microfluidic device [38] was adapted for imaging to create a well-controlled, no-flow environment. Before the experiments, devices were filled with a 1% MC solution and kept for at least 2 hours in an incubator at 38.5°C with 5% of CO_2_. Sperm suspension was then seeded onto the 2 mm access hole of the device and sperm were allowed to swim inside the device. The microfluidic chamber is 100 *μ*M deep, and we captured videos close to the lower surface by a scientific CMOS camera (ANDOR Neo for Sample A (fresh) and Zyla for Sample B (frozen)) at their respective highest frame rates (135-165 frames per second), in conjunction with an inverted microscope with phase contrast and NIS-Elements software. Given an estimated 20 *μ*m depth of the focal plane, only vague shadows of sperm swimming close to the upper surface were seen. The devices were maintained at 38.5°C during the experiments.

### 4.2 Extraction and parameterisation of flagella

#### 4.2.1 Tracking and selection

Videomicroscopy data is processed using the TrackMate plugin included in the popular software package Fiji [26, 39, 40], with TrackMate automatically tracking the locations of the spermatozoa between frames by identifying their cell bodies. Individual swimmers are then isolated using bespoke Fiji macros, utilising automatic local thresholding to segment individual swimmers into binary masks. Each swimmer mask is processed using the fully automated scheme of Walker et al. [18], with flagella being identified by their approximately consistent visible width. The resulting digital representations of the flagella, validated by eye and consisting of pixel locations of the flagellum in each frame, are screened to accept only those where the observable flagellum length, projected onto the focal plane of the microscope, varied by less than 10% of the maximum over multiple beating periods. Of these accepted individuals, frames containing digitised flagella of captured length greater than this 90% threshold were truncated in space to enforce uniformity whilst avoiding the need for extrapolation. This truncated flagellar length represents the true flagellar length to within approximately 10%. Reliably reporting the true flagellar length is precluded by difficulties in imaging the most-distal region of the flagellum, an issue common in the microscopy of flagellates though here limited to approximately the distal 10%.

#### 4.2.2 Spatial and temporal smoothing

With the data by construction reporting spatiotemporal information for approximately 90% of the visible flagellum for each swimmer, pixel locations are identified with Cartesian *xy* coordinates, translated and rotated so that the proximal flagellar end is both at the origin and aligned along the horizontal axis (1, 0), as illustrated in fig. 1 (lower left), hence quantifying flagellar motion relative to the swimmer body. Each flagellum is parameterised by arclength and time, and smoothing in both space and time performed using smoothing splines and Gaussian convolutions, respectively, in the software package MATLAB^®^. Owing to infrequent erroneous results of the flagellar identification and tracking process, a low proportion (less than 10%) of frames corresponding to each individual swimmer are omitted from temporal smoothing due to such errors. Instead, for these rare frames, flagellum data is linearly interpolated from the preceeding and following frames, and we note in particular that the frame rates are sufficiently high so as to preclude significant interpolation errors. All smoothing and interpolation is validated by visual comparison with the unsmoothed data.

#### 4.2.3 Determining the beating period

Whilst existing CASA implementations are able to approximate the beating period of spermatozoa from the oscillations of the body over time, as first suggested by Schoëvaërt-Brossault [41] and Serres et al. [42] and more recently conducted via Fourier analysis, we opt to determine the period of the flagellar beat using the captured flagellar kinematics. The recent work of Gallagher et al. [14] also based their estimate of the period on captured flagellar data, though utilised the Fourier spectra of points along the flagellum to identify a dominant frequency, implicitly requiring that the beating period is comprised of a single sinusoidal mode. Here we avoid this requirement by computing the period using the autocorrelation of a material point, which may be thought of as tracking the location of this point over time and identifying the period after which its trajectory is most self-similar. Performing this computation using multiple points along the flagellum, the minimal beating period of each flagellar waveform may be automatically identified, which we denote by *T*, with results verified by eye. Indeed, we note that sampling each flagellum only at its midpoint was sufficient in this case to reliably determine the beating period.

#### 4.2.4 Normalisation and discretisation

The results of spatial and temporal smoothing are natural Cartesian representations of each flagellar waveform, denoted 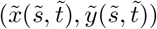, where for each individual the arclength, 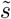, ranges from zero to the captured flagellum length, *L*, and 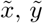 are relative to the sperm reference frame, as detailed in the section on data smoothing and as used in the lower left panel of fig. 1. The time, 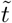, encompasses the timestamps of the captured frames containing this flagellum, with the first frame occuring at 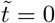 without loss of generality, and the timestamp of the final frame being denoted *T_f_*. In order to facilitate comparison between individuals, each waveform is normalised in space by the length of the flagellum, so that the rescaled arclength, 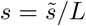, lies between zero and one for all of the swimmers. A priori it is unclear whether this rescaling of arclength is appropriate, particularly if there were to be significant variation in length over the population. However, as demonstrated in fig. 3, the distribution of measured flagellum length is in fact tightly clustered about its mean, thus this rescaling in space represents a minimal transformation of the original data, and as such is used throughout this work. The waveforms are then discretised at 1000 points in space, capturing at each instant in time the locations of material points equally spaced along the flagellum at fixed arclengths *s_i_* = (*i* − 1)/999, *i* = 1,…, 1000.

Similarly, the temporal dynamics of each swimmer are rescaled by their beating period, with rescaled time being denoted 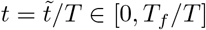 and the fully-normalised waveform written as (*x*(*s, t*), *y*(*s,t*)). Before being discretised in time, each waveform is temporally truncated such that it represents only a single beat of the captured flagellum, i.e. it is restricted to some interval *t* ∈ [*t*^⋆^, *t*^⋆^ + 1), noting that the period of the rescaled beat is one. The offset *t*^⋆^ is found for each swimmer by comparing the locations of the flagellum midpoint, that corresponding to *s*_500_, at times *t* and *t* + 1 for all valid *t* ∈ [0, *T_f_*/*T*], selecting *t*^⋆^ so as to minimise the Euclidean distance between the locations of the material point at each instant. The temporally-truncated waveforms are then discretised into 100 equispaced timepoints, and all waveforms are aligned in phase with one another by prescribing the location of a fixed phase in the periodic behaviour, with this simple method of phase alignment being verified a posteriori.

#### 4.2.5 Representations of the flagellum

Above we have described normalised waveforms using a natural representation of the rescaled flagellar beat in spatial Cartesian coordinates, (*x*(*s, t*), *y*(*s,t*)), where *s* and *t* denote normalised arclength and time parameters. However, in some cases it is more convenient to consider each beat in terms of its tangent angle parameterisation, for example when generating synthetic beating patterns. This tangent angle parameterisation, here denoted by *θ*(*s, t*), is defined by the nonlinear relation

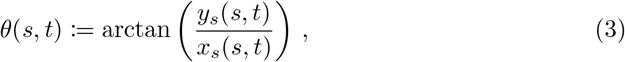

where here *y_s_*(*s, t*) and *x_s_*(*s, t*) denote the derivatives of *y*(*s, t*) and *x*(*s, t*) with respect to normalised arclength *s*, and arctan is the four-quadrant inverse tangent function (e.g. the function `atan2` in MATLAB^®^). This relation is readily inverted to allow the recovery of the Cartesian beating pattern from the tangent angle in the reference frame of the spermatozoon, as described in the section on flagellum smoothing and illustrated in fig. 1 (lower left):

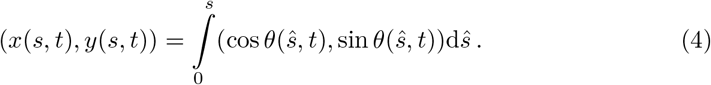

In practice these derivatives, integrals, and function evaluations must be performed numerically, and we observe negligible cumulative error when transforming between parameterisations using 1000 material points to represent each flagellum.

### 4.3 Waveform analysis

#### 4.3.1 Principal component analysis

A standard tool for dimensionality reduction in various contexts, principal component analysis (PCA) may be used to identify the most significant constituents of the flagellar beat. Given multiple observations of a quantity, which in our case will either be *xy* coordinates or the accompanying tangent angle parameterisation of material points along the flagellum, PCA outputs time-independent components or *modes*, along with time-dependent *coefficients*, with the coefficients for the *i*th mode here being denoted *c_i_*. These PCA modes and coefficients are given in descending order of their contributions to the variance of the observed quantities, with their contributions quantified by *weights* that sum to unity. For a technical description of principal component analysis, we direct the interested reader to the relevant work of Werner et al. [20]. In the study presented here, at a timepoint *t_j_* our observations will be of the form (*x*_1_, *y*_1_, …, *x*_1000_, *y*_1000_) or (*θ*_1_, …, *θ*_1000_), corresponding to Cartesian and tangent angle representations of the waveform, respectively, where *x_i_* denotes *x*(*s_i_*, *t_j_*) for the *i*th equispaced arclength *s_i_* at the *k*th timepoint *t_k_*, with analogous definitions for *y_i_* and *θ_i_*.

PCA in the context of flagellar analysis is often performed on a set of observations taken from an individual cell, yielding for that individual a set of modes, coefficients and weights. Whilst this may provide insight into the constituent components of a particular waveform, its utility in comparing the beats of different swimmers is limited to consideration of the PCA modes, with the coefficients not being comparable due to their ties to their respective, and different, modes. Further, as modes corresponding to individual swimmers can represent different contributions to the overall beating, given by their individual weights, comparisons between the modes of different swimmers are themselves very limited in scope. Thus, in this work we apply principal component analysis to all the swimmers in a given sample at once, combining the observations of the motion of each material point across the population. This enables us to comment not only on individual variation, but on the quantitative variation of waveforms across the population, with the PCA coefficients now comparable between swimmers as they are each with respect to a single common set of modes. Magnitudes of contributions of different modes are compared via the relevant coefficients, with the modes normalised such as to render this comparison meaningful.

#### 4.3.2 Maximal tip curvature (MTC)

Given a tangent angle parameterisation *θ*(*s, t*) of a flagellar waveform, as defined by (3), the signed curvature, *κ*, is given by *κ* := *θ_s_*(*s, t*), where again the subscript denotes a derivative with respect to normalised arclength. Numerically computing this curvature over the distal 10% of the captured flagellum, we take the maximum of its absolute value over this distal flagellar portion and the entire beating period, and report this quantity as the maximal tip curvature.

### 4.4 Statistical tests

Distributions of the first Cartesian PCA coefficient are compared via linear least squares fitting of the coefficient for each individual swimmer, where the linear model to be fitted is of the form

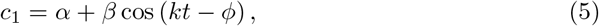

informed by the approximately sinusoidal form of the average coefficient shown in fig. 4. Here, *α* and *β* are constant in arclength and time, but vary between sperm. In addition, *t* again represents normalised time, and wavenumber and phase parameters *k* and *ϕ* are first approximated by non-linear least squares fitting of the mean coefficient, 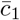, (see fig. 2, right panels). This is implemented in MATLAB^®^, in particular keeping both *k* and *ϕ* fixed throughout the linear fits. Differences between fits are quantified by standard two-tailed two-sample Kolmogorov-Smirnov (KS) tests performed on the distributions of the parameters *α* and *β*, with the conclusion of an overall significant difference drawn only if both tests are significant at the 5% level. Refinements are possible on larger sample sizes using the joint distribution of fit parameters, though are not implemented in this work. All other statistical tests performed are either KS tests or Wilcoxon rank-sum tests, each at the 5% significance level.

## 5 Data Access

The data used and generated in this paper is freely available from the University of Oxford Research Archive (ORA) via the link http://dx.doi.org/xxx/xxx.

## 6 Acknowledgements

The authors acknowledge Prof. Adriana Dawes for helpful discussions on the subject of statistical testing. Experimental data acquisition was supported by National Science Foundation (HRD 1665004 to C.K.T) and National Institutes of Health (R15HD095411 to C.K.T). B.J.W. is supported by the UK Engineering and Physical Sciences Research Council (EPSRC), grant EP/N509711/1. K.I. is supported by JSPS-KAKENHI for Young Researchers (18K13456) and JST, PRESTO Grant Number JPMJPR1921, Japan. This work made use of the Cornell Center for Materials Research Shared Facilities which are supported through the NSF MRSEC program (DMR-1719875).

## 7 Competing Interests

The authors declare no competing interests.

